# Boundary Bypass Activity in the *Abdominal-B* Region of the *Drosophila* Bithorax Complex is Position Dependent and Regulated

**DOI:** 10.1101/2023.06.06.543971

**Authors:** Olga Kyrchanova, Airat Ibragimov, Nikolay Postika, Pavel Georgiev, Paul Schedl

## Abstract

Expression of *Abdominal-B* (*Abd-B*) in abdominal segments A5 *–* A8 is controlled by four regulatory domains, *iab-5 – iab-8*. Each domain has an initiator element (which sets the activity state), elements that maintain this state and tissue-specific enhancers. To ensure their functional autonomy, each domain is bracketed by boundary elements (*Mcp*, *Fab-7*, *Fab-7* and *Fab-8*). In addition to blocking crosstalk between adjacent regulatory domains, the *Fab* boundaries must also have bypass activity so the relevant regulatory domains can “jump over” intervening boundaries and activate the *Abd-B* promoter. In the studies reported here we have investigated the parameters governing bypass activity. We find that the bypass elements in the *Fab-7* and *Fab-8* boundaries must be located in the regulatory domain that is responsible for driving *Abd-B* expression. We suggest that bypass activity may also be subject to regulation.

**Summary Statement:** Boundaries separating *Abd-B* regulatory domains block crosstalk between domains and mediate their interactions with *Abd-B*. The latter function is location but not orientation dependent.

## INTRODUCTION

The mechanisms regulating gene expression in multicellular eukaryotes are intimately connected to the 3D organization of the genome. Chromosomes are subdivided into a series of looped domains called TADs (topologically associated domains) by special elements called boundaries or insulators [1–6]. In mammals, the main protein implicated in boundary function is the multi-zinc finger protein CTCF and in ChIP experiments it localizes to the endpoints of many mammalian TADs [7]. By way of contrast, in *Drosophila* more than a dozen proteins including not only CTCF (dCTCF), but also other several other multi-zinc finger proteins (Pita, M1BP, Zipic, Zw5, and Su(Hw)) have been shown to have boundary function and these proteins ChIP to sequences that define the endpoints of fly TADs [8–12].

In addition to determining the 3D organization of chromosomes, boundary elements have genetic functions. When interposed between enhancers/silencers and genes they can block regulatory interactions [13–19]. As a consequence of this blocking activity, when chromosomal segments are flanked by boundary elements, they define units of independent genetic activity. In this case, enhancers/silencers and genes residing within the same TAD engage in regulatory interactions, while cross-TAD interactions are suppressed [20,21]. However, there are instances in which regulatory interactions must take place between enhancers/silencers in one insulated domain and genes located in another insulated domain. For example, the distant regulatory elements driving expression of *HoxD13-HoxD10* gene cluster are separated from their target genes by multiple sites for the CTCF boundary factor [22]. Similarly, the murine Sonic Hedgehog gene is regulated by multiple enhancers that spread over nearly a Mb (megabase) and span several TADs [23,24]. In both of these cases, the enhancers must bypass one or more boundary elements in order to interact with their target promoters. In addition to reaching over large distances and across multiple TADs, there must be mechanisms in place to ensure specificity, otherwise the enhancers could interact with the wrong genes. As is the case for these two vertebrate genes, the parasegment specific regulatory domains in the *Drosophila melanogaster* Bithorax complex must also be able to bypass one or more intervening boundary element in order regulate their gene targets. However, while little is currently known about the mechanisms or elements involved in mediating cross-TAD regulatory interactions in vertebrates, the *cis-*acting elements and the *trans-*acting factors responsible for boundary bypass have been identified in flies.

The three BX-C homeotic genes, *Ultrabithorax* (*Ubx*), *abdominal-A* (*abd-A*) and *Abdominal-B* (*Abd-B*) determine parasegment (segment) identity in the posterior 2/3rds of the fly, from parasegment PS5 to PS14 [25–30]. Specification of PS identity in PS5-PS13 depends upon nine parasegment specific regulatory domains that are responsible for directing the appropriate temporal and spatial pattern of expression of one of the homeotic genes (Fig 1A). The *Ubx* gene functions in the specification of PS5 (segment T3 in the adult cuticle) and PS6 (segment A1) and it is expression in these two parasegments is controlled by the *abx/bx* and *pbx/bxd* domains, respectively. Three domains, *iab-2*, *iab-3* and *iab-4* control *abd-A* expression in PS7(A2), PS8(A3) and PS9(A4). Finally, the *iab-5*, *iab-6*, *iab-7* and *iab-8* domains regulate *Abd-B* expression in PS10(A5), PS11(A6), PS12(A7), and PS13(A8) (Fig. 1A).

**Fig. 1.**
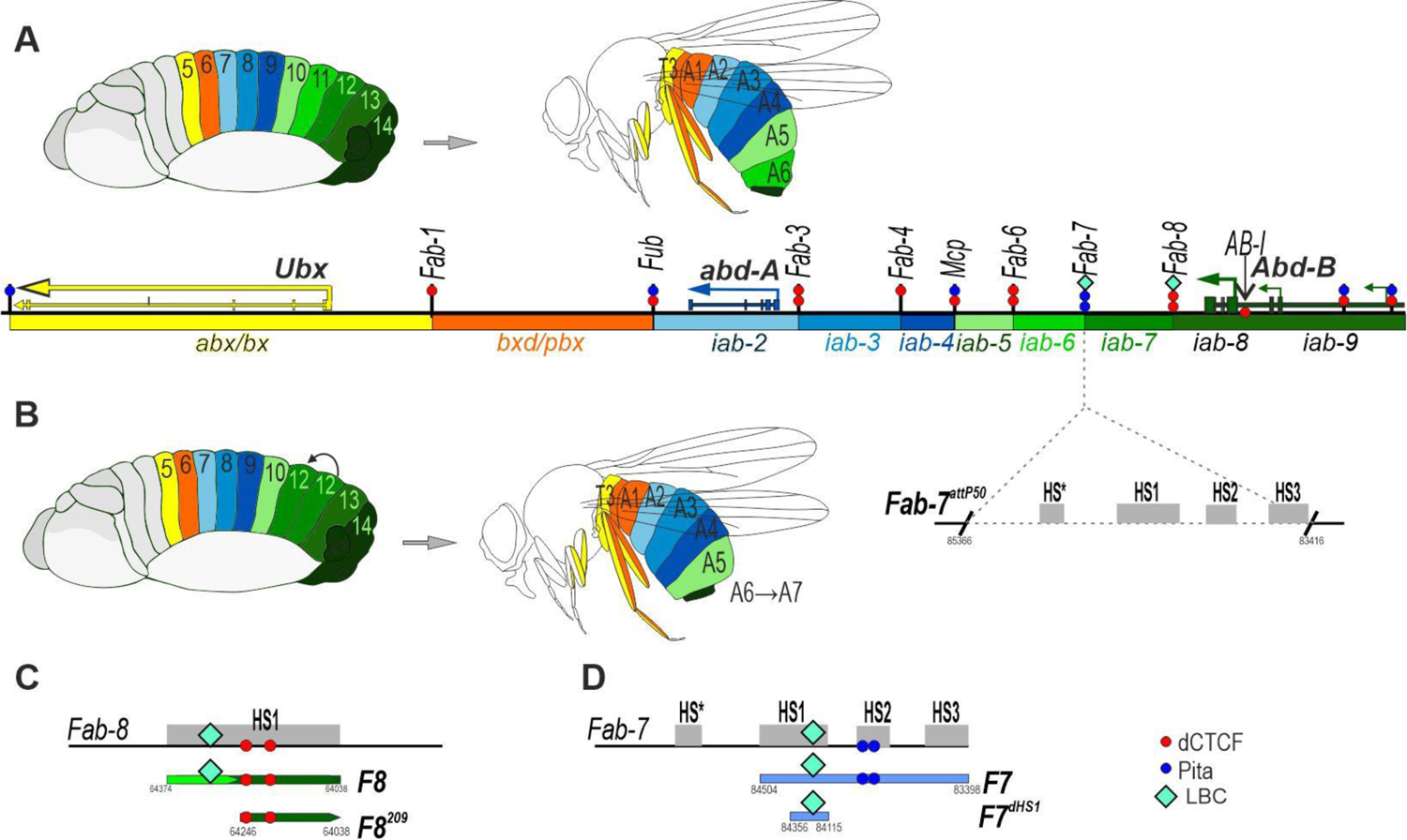
A) Map of the *Ubx, abd-A* and *Abd-B* regions of the *Drosophila melanogaster* BX-C. The *Ubx* gene is regulated by *abx/bx* and *bxd/pbx* domains (marked with yellow and orange) in parasegments PS5 and PS6 respectively (which correspond approximately to segments T3 and A1 in adults). Three regulatory domains, *iab-2*, *iab-3* and *iab-4* (shades of blue), control *abd-A* expression in PS7(A2), PS8(A3), PS4(A4) respectively. *Abd-B* expression in PS10(A5), PS11(A6), PS12(A7) and PS13(A8) is controlled by *iab-5*, *iab-6*, *iab-7* and *iab-8* (shades of green) respectively. The embryo and adult segments are indicated using the same color code as the *iab* domain that is required for their specification. The black lines with colored circles mark chromatin boundaries: *Fab*-*1*, *Fub*, *Fab-3, Fab-4*, *Mcp*, *Fab-6*, *Fab-7*, and *Fab-8*. The red circles indicate the number of CTCF binding sites in each boundary, and the blue – the number of Pita sites. LBC – as cyan. **(B) Deletion of the *Fab-7* boundary.** When the *Fab-7* boundary is deleted, *iab-6* and *iab-7* fuse into one domain. As a result, PS11/A6 is transformed into a copy of PS12/A7. The *Fab-7^attP50^* platform, in which four hypersensitive regions, HS*, HS1, HS2, and HS3 (marked with gray boxes) are deleted, is indicated by broken black lines. **Maps of the (C) *Fab-7* and (D) *Fab-8* fragments**. Maps of fragments that were used for replacements. The *Fab-8* fragments are indicated as follows: the bypass element is indicated by the light green line, the insulator element by the dark green line. The coordinates are according to the complete sequence of BX-C in the SEQ89E numbering [71]. The *Fab-7* fragments are shown as a light blue line.

Each regulatory domain has an initiator element that sets the activity state of the domain, *on* or *off* early in embryogenesis [27,28,31]. Initiators respond to the maternal, gap and pair-rule gene products that subdivide blastoderm stage embryos along the antero-posterior axis into 14 parasegments [32–38]. For example, in PS11(A5), the *iab-5* initiator turns *on* the *iab-5* domain, while the adjacent *iab-6* and other more distal (relative to centromere) domains remain in the *off* state. In PS11(A6), the initiator in *iab-6* turns the domain *on*. While *iab-5* is also active in PS11, *iab-7* and *iab-8* are *off*. The gene products responsible for setting the activity state of the BX-C domains disappear during gastrulation and different mechanisms are used to remember the activity state during the remainder of development. The *off* state is maintained by Polycomb group (PcG) silencing, while remembering the *on* state requires proteins in the trithorax (Trx) group [28,39– 41]. The regulatory domains also contain a series of tissue and stage specific enhancers which are responsible for driving the expression of their cognate homeotic gene in a pattern appropriate for the differentiation of the parasegment (segment) they specify [31,38].

In order to specify PS identity, regulatory domains in BX-C must be functionally autonomous. Functionally autonomy is conferred by the boundary elements that flank each regulatory domain and are responsible for blocking crosstalk between regulatory elements in adjacent domains [13,14,17,19,42–46]. The genetic and developmental roles of the four boundaries in the *Abd-B* region of the complex, *Mcp*, *Fab-6*, *Fab-7* and *Fab-8*, are among the most thoroughly studied and understood in multicellular eukaryotes. The centromere proximal boundary *Mcp* is located between *iab-4* and *iab-5* and it marks the border separating the regulatory domains for *abd-A* and *Abd-B*. *Fab-6* is located between *iab-5* and *iab-6*, *Fab-7* between *iab-6* and *iab-7* and *Fab-8* between *iab-7* and *iab-8* (Fig. 1A). Deletion of one of these boundaries has a profound effect on development, resulting in a gain-of-function (GOF) transformation in parasegment identity. For example, when *Fab-7* is deleted, the initiation element in *iab-6* ectopically activates the *iab-7* domain in PS11(A6) (Fig. 1B). As a result, *iab-7* drives *Abd-B* expression not only in PS12/A7 but also in PS11/A6, transforming PS11/A6 into a copy of PS12/A7 [45]. Similar GOF transformations are observed for deletions of *Mcp*, *Fab-6* and *Fab-8* [13,35,43,47].

In addition to blocking crosstalk between adjacent regulatory domains, *Fab-6*, *Fab-7* and *Fab-8*, but not *Mcp*, must also support long-distance regulation so that enhancers located in the *iab-5*, *iab-6* and *iab-7* domains can bypass the intervening boundaries and communicate with the *Abd-B* promoter [48–50]. The requirement for bypass activity is most clearly evident when the *Fab-7* boundary is replaced by a heterologous fly boundary or multimerized binding sites for zinc finger proteins like dCTCF, Pita or Su(Hw). These foreign elements are typically able to prevent crosstalk between *iab-6* and *iab-7* and rescue the GOF transformation of PS11/A6 into PS12/A7. However, because they are unable to mediate boundary bypass, the *iab-6* domain cannot activate *Abd-B* in PS11/A6 cells [49,51,52]. As a consequence, PS11/A6 is transformed towards a PS10/A5 identity. Not surprisingly given its location separating the regulatory domains for *abd-A* and *Abd-B*, the *Mcp* boundary is able to block crosstalk, but cannot support bypass [50].

In previous studies we found that *Fab-7* and *Fab-8* have sub-elements (Fig. 1C,D) that primarily (but not exclusively) function either as insulators and block crosstalk between adjacent domains or as bypass elements that mediate long distance regulatory interactions [49,53,54]. In the case of *Fab-8*, blocking activity is conferred by a 209 bp centromere distal fragment that contains two binding sites for CTCF. Bypass activity is conferred by a proximal 165 bp fragment [54] (Fig. 1C). In nuclear extracts this fragment is shifted by a large multiprotein complex called LBC that is thought to contain GAF, Mod(mdg4) and e(y)2, while ChIP experiments indicate that it is also bound by CLAMP [55]. The *Fab-7* boundary spans four hypersensitive sites, HS*, HS1, HS2 and HS3; however, it is possible to reconstitute a fully functional boundary (blocking and bypass) by combining two 200 bp fragments corresponding to the distal half of HS1, dHS1, and HS3 [49,53] (Fig. 1D). HS3 not only provides boundary activity, it also functions as a Polycomb Response Element, PRE [56,57]. Like the 165 bp *Fab-8* fragment, dHS1 is shifted by the LBC in nuclear extracts [54,55]. While dHS1 is necessary for blocking crosstalk, it also has bypass activity. *Fab-7* boundary activity can be fully reconstituted by combining dHS1 with multimerized bindings sites for the zinc finger proteins Pita, Su(Hw) or dCTCF [49]. For example, a 5× multimer of Pita (Pita^×5^) blocks crosstalk but does not support bypass; however, when combined with dHS1 (*dHS1+Pita^×5^*) the artificial boundary fully rescues the *Fab-7* deletion. Moreover, it would appear that bypass activity is an active process as the *dHS1+Pita^×5^*combination induces a GOF transformation when used to replace the *Mcp* boundary [50]. As noted above, *Mcp* marks the border between the *abd-A* and *Abd-B* regulatory domains. When *dHS1+Pita^×5^*is substituted for *Mcp* it induces the *abd-A* regulatory domain *iab-4* to inappropriately activate *Abd-B* expression in PS9/A4.

To better understand the functional requirements for bypass activity we used the *Fab-7* deletion, *Fab-7^attP50^* [55] to manipulate elements conferring blocking and bypass activity. Our experiments indicate that the order of the bypass and blocking elements is important for full bypass activity. The bypass element must flank the domain, in this case *iab-6*, which requires bypass activity to activate *Abd-B* while block element flanks the adjacent, inactive domain, *iab-7*. When the order is reversed, blocking but not bypass activity is observed.

## RESULTS

### Orientation or order?

In previous experiments we replaced the *Fab-7* boundary with the neighboring *Fab-8* boundary [48]. When a minimal 337 bp *Fab-8* element, F8 (Fig. 1C; Fig. 2A), was inserted in the forward orientation (the same as the endogenous *Fab-8*) it fully substituted for *Fab-7*: it blocked crosstalk between *iab-6* and *iab-7* and supported bypass activity, enabling *iab-6* to regulate *Abd-B* expression in PS10(A5) (Fig. 2B and 2C). On the hand, when F8 was inserted in the reverse orientation, F8^R^ (Fig. 2A), it blocked crosstalk, but did not fully support bypass (Fig. 2B and 2C). In this case, PS11(A6) was partially transformed towards PS10(A5) because *iab-6* is unable to properly regulate *Abd-B* in PS11. As can be seen in Fig. 2B, the morphology of the A6 segment in the F8 (forward) replacement resembles *wild type* (*wt*). In darkfield images, the trichome hairs on the A6 tergite are restricted to the anterior and ventral margins, while the sternite has a characteristic banana shape. In contrast, in the F8^R^ replacement, there are ectopic trichome hairs on the dorsal side of the A6 tergite, while the A6 sternite has an abnormal shape (in between the banana shape of A6 and the quadrilateral shape of A5) and has several bristles. Consistent with these phenotypic effects in adult males, expression of Abd-B in PS11 in the embryonic CNS is reduced compared to *wt* and absent in PS10 (Fig. 2C, Fig. S1 shows Abd-B plus Engrailed to indicate parasegment borders).

**Fig. 2.**
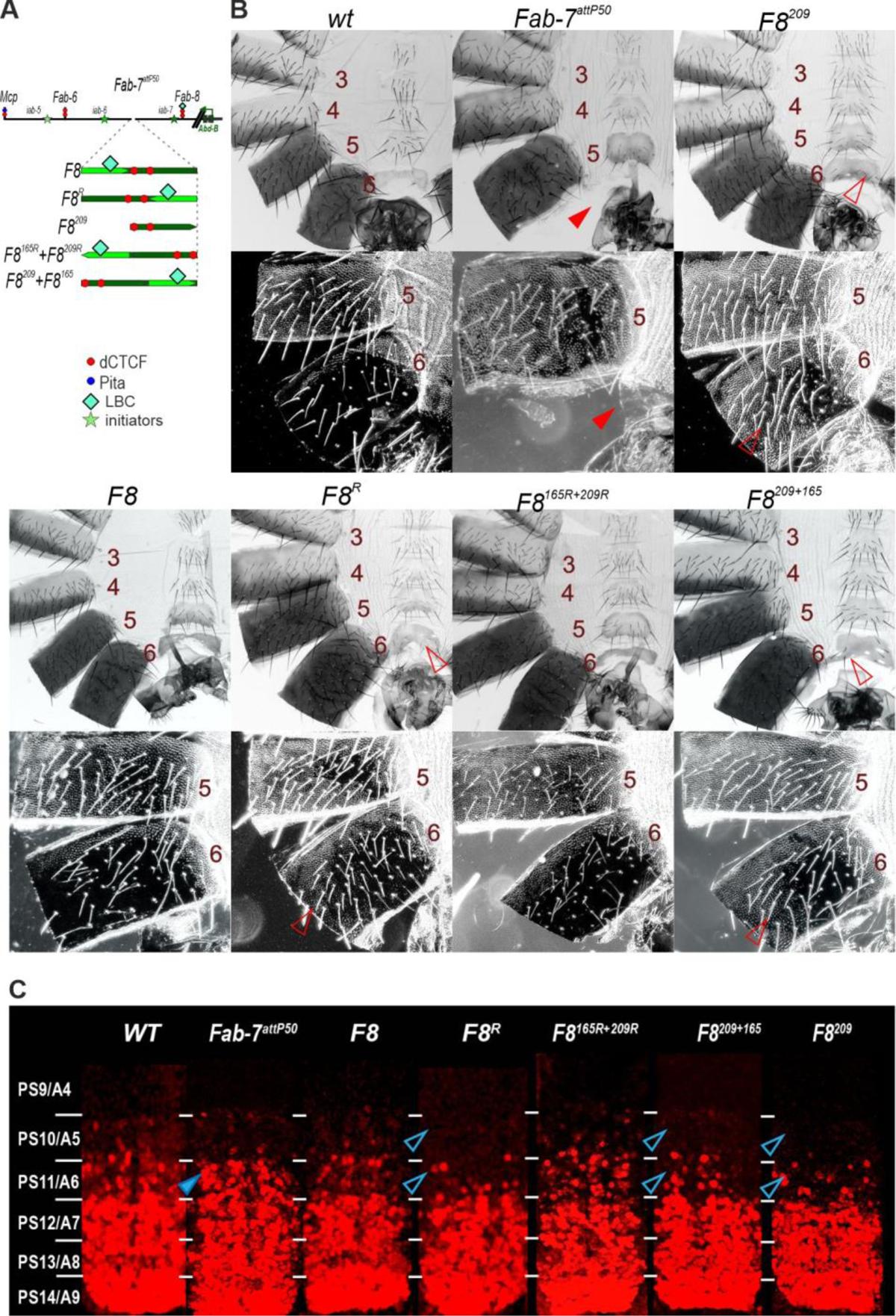
The *Fab-8* bypass element (F8^165^) must be located adjacent to *iab-6* in order to overcome the blocking activity of the *Fab-8* insulator (F8^209^). (A) Scheme of the *Abd-B* regulatory domains and fragments used for *Fab-7* replacements. All other designation as in Fig. 1. **(B)** Morphology of the male abdominal segments (numbered) in wild type (*wt*) and in F8, F8^R^, F8^209^, F8^165R^+F8^209R^, F8^209^+F8^165^. Trichomes on the A5 and A6 tergites are shown in dark field. In *wt* the A6 sternite has a banana shape and is devoid of bristles, while the A5 sternite has a quadrilateral shape and is covered in bristles. The A6 tergite has trichomes along the anterior and ventral edges, while the entire A5 tergite is typically covered in trichomes with small internal patch that lack trichomes. The filled red arrowheads show morphological features indicative of GOF transformations, the empty arrowheads – LOF transformations. **(C)** *Abd-B* expression in the CNS of *wt* and the *Fab-7* replacement lines. Each panel shows an image of the CNS of stage 14 embryos stained with antibodies to Abd-B (red). White horizontal bars delimit parasegment boundaries. Parasegments are numbered from 9 to 14 on the right side of the panels; approximate positions of segments are shown on the left side of the *wt* panel and marked 4 to 8. The *wt* expression pattern of *Abd-B* in the embryonic CNS is characterized by a stepwise gradient of increasing protein level from PS10 to PS14. Staining for Engrailed was used to define the boundaries of the parasegments (Fig S1).

Since pairing interactions between fly boundaries are typically orientation dependent [15,58], we previously suggested that the loss of bypass activity in the F8^R^ replacement was likely a consequence of an altered loop topology. However, as noted above, *Fab-8* blocking is mostly dependent on a centromere distal 209 bp fragment, F8^209^, (Fig. 2A), while a proximal 165 bp fragment, F8^165^ confers bypass activity [54]. The fact that blocking and bypass are largely mediated by two distinct DNA elements raised the possibility that it is their order rather than their orientation that is critical. To test this possibility, we generated two replacements. In the first, we inverted the F8^165^ and F8^209^ fragments, but kept their order with respect to *Abd-B* the same as the endogenous *Fab-8* boundary (Fig. 2A: F8^165R^+F8^209R^). In the second we kept the endogenous orientation of F8^165^ and F8^209^, but reversed their order with respect to *Abd-B* (Fig. 2A: F8^209^+F8^165^). In this case, F8^209^ boarders the *iab-6* domain, while F6^165^ is adjacent to the *iab-7* domain.

Fig. 2B shows that it is the order of the bypass and blocking fragments, not their orientation that is important. The F8^165R^+F8^209R^ replacement has a *wt* phenotype. The A6 sternite has the characteristic banana shape, while the trichome hairs on the tergite are restricted to the anterior and ventral edges. In contrast, F8^209^+ F8^165^ has blocking activity, but does not fully support bypass: the A6 sternite has an almost quadrilateral shape like in A5, while there are ectopic trichome hairs on the tergite. The phenotypic effects seen in the adult male A6 cuticle are recapitulated in the pattern of *Abd-B* expression in the embryonic CNS. Whereas F8^165R^+F8^209R^ resembles *wt* or F8, the pattern of *Abd-B* expression in the CNS in F8^209^+F8^165^ is closer to that of F8^R^ (Fig. 2C, Fig. S1).

### The blocking function F8^209^ can interfere with *Fab-7* dependent bypass

It seemed possible that the element conferring bypass in *Fab-7* replacements might need to be next to the *iab-6* domain in order to mediate interactions between *iab-6* and *Abd-B*. To explore this possibility, we generated a composite boundary, F8^209^+F7, in which F8^209^ is next to the *iab-6* domain (Fig. 3). We reasoned that since both F8^209^ and *Fab-7* are in their normal forward orientations, the orientation of pairing interactions with elements (e.g., AB-I (Fig. 1A) [59]) upstream of the *Abd-B* promoter should not be affected. On the other hand, if the order of the elements with bypass and blocking activity is important, then the bypass activity of *Fab-7* should be disrupted.

**Fig. 3.**
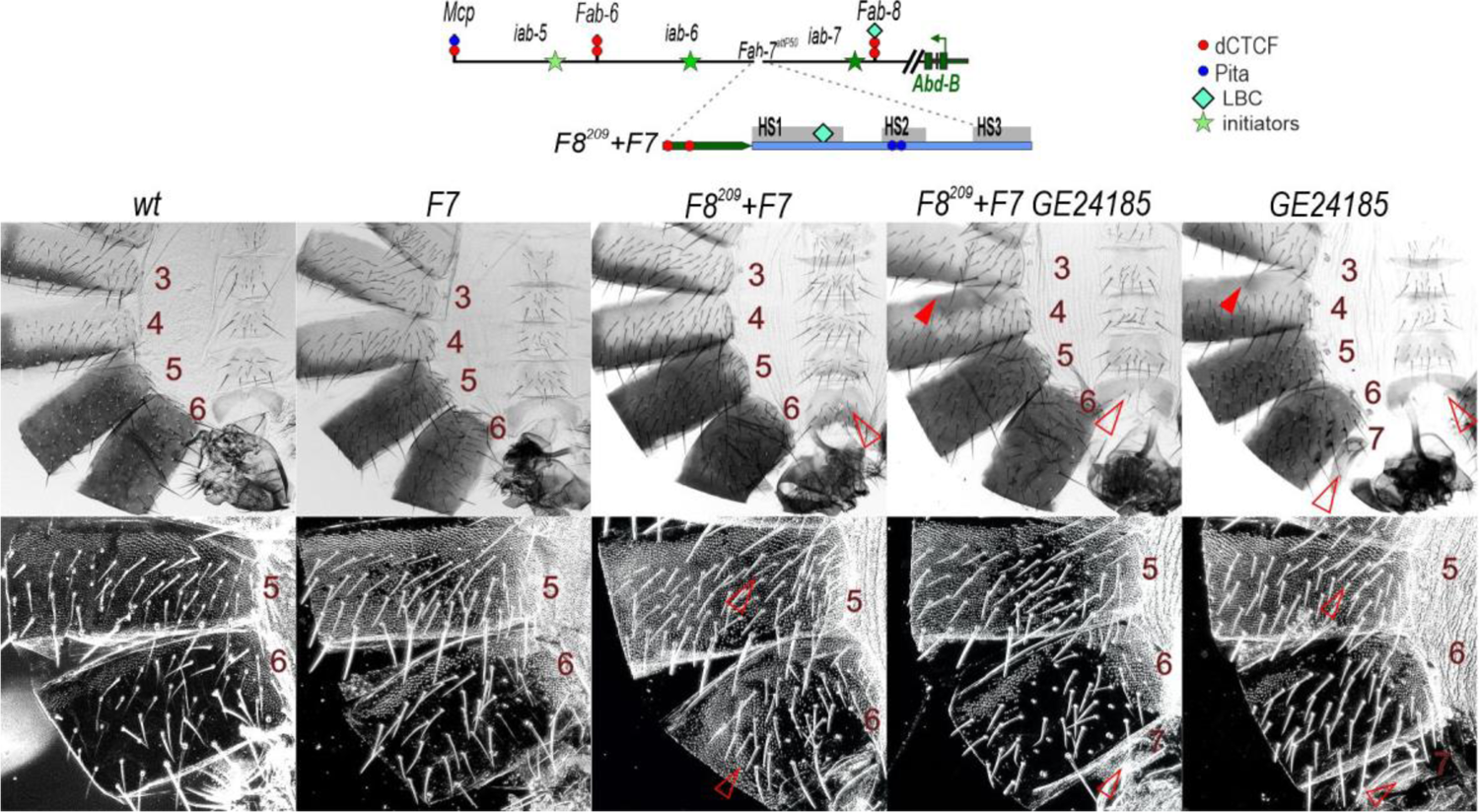
Testing the ability of F8^209^ to block the *Fab-7* dependent bypass. (A) Map of the *Abd-B* regulatory region and F8^209^+F7 fragment used for *Fab-7* replacements. All designation as in Fig. 1 (B) Bright field (top) and dark field (bottom) images of cuticles prepared from *wt*, F8^209^, F8^209^+F7, F8^209^+F7+GE24185, GE24185, where GE24185 is null *dCTCF* allele in GE24185/GE24185 flies[60]. The empty red arrowheads point to signs of LOF transformations, which are correlated with the loss/lack of bypass functions of the tested DNA fragments. The filled red arrowheads show morphological features indicative of GOF transformations.

F8^209^ alone suppresses the GOF transformation of the starting *Fab-7* deletion platform *Fab-7^attP50^*; however, it does not fully support bypass, resulting in a loss of function (LOF) transformation of PS11 (A6) towards PS10 (A5) [48]. As shown in Fig. 3, the A6 tergite has ectopic trichome hairs, while the sternite is misshapen and has bristles. When F8^209^ is placed in front of a fully functional *Fab-7* boundary it has the same effect; it interferes with the bypass activity of *Fab-7*. Much of the A6 tergite is covered in trichome hairs, while the LOF phenotype of the sternite becomes even more pronounced — it has a quadrilateral shape and is covered in bristles (Fig. 3).

*Fab-8* blocking activity requires the two dCTCF binding sites in the F8^209^ fragment [48]. If F8^209^ blocking is responsible for the loss of bypass activity in the F8^209^+F7 combination, one would expect that this defect might be partially ameliorated by introducing a mutation in the fly *ctcf* gene (GE24185) [60,61]. The GE24185 mutation in a *wt Fab-7* background disrupts *Abd-B* expression (Fig. 3). As dCTCF is important for *Mcp* boundary activity, some GE24185 males have ectopic pigmentation in A4. This is also seen in the F8^209^+F7 GE24185 combination. There are also weak LOF transformations in A6 and A7. The A6 sternite has a *wt* banana shape, but has several bristles. While these phenotypes are evident in F8^209^+F7 GE24185 males, the bypass defects induced by F8^209^ are partially rescued. Instead of a quadrilateral shape, the A6 sternite has a banana shape, while the tergite is only partially covered in trichome hairs (much like GE24185 alone).

### The *gypsy su(Hw)* insulator disrupts *Fab-7* bypass activity

The results in the previous sections suggest that relative order of elements with bypass and blocking activity is critical for activation of *Abd-B* by the *iab-6* enhancers. If this is the case, then the bypass activity of F7 should be disrupted if entirely heterologous boundaries are placed between it and the *iab-6* domain. To test this prediction, we used the *gypsy* Su(Hw) insulator (*gy*) in combination with F7 (Fig. 4A). Previous studies by Hogga et al. [51] showed that when *Fab-7* is replaced by *gy* it blocks *iab-6:iab-7* crosstalk but does not support bypass (Fig. 4B). However, the effects of the *gy* replacement on the morphology of the adult cuticle and *Abd-B* expression in the embryonic CNS are somewhat different from that described above. In the adult male cuticle, the transformation of A6 into A5 is more complete than in either F8^R^ and F8^208^. The A6 tergite is covered in trichome hairs, while the sternite has a quadrilateral shape just like A5 and is covered in bristles. In addition, the morphology of the A5 segment has features indicative of a transformation towards an A4 identity (Fig. 4B). There are patches of cuticle in the A5 tergite that lack pigmentation, while the trichome hairs are densely packed like the A4 tergite. On the other hand, Abd-B expression in PS11 in the embryonic CNS is elevated compared to *wt*, indicative of a GOF rather than an LOF transformation in parasegment identity. A similar result was reported by Hogga et al [51].

**Fig. 4.**
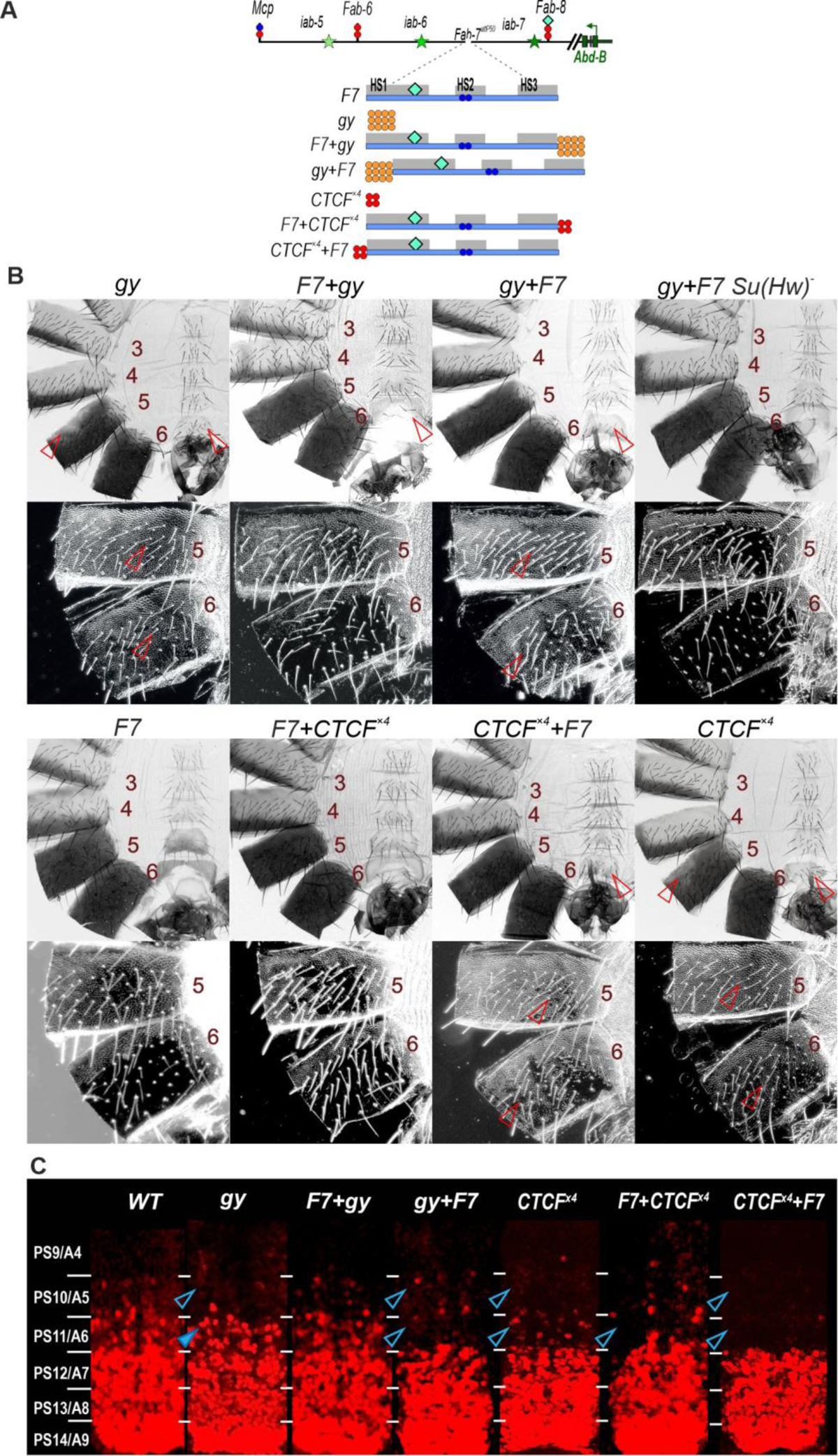
The *gypsy* (Su(Hw)) and CTCF^×4^ insulators block *iab-6* enhancers from regulating *Abd-B* when placed next to the *iab-6* domain. (A) Map of the *Abd-B* regulatory region and fragments used for *Fab-7* replacements. All designation as in Fig. 1 and 2. *gy* is shown as 12 orange circles reflecting 12 binding sites for Su(Hw) in insulator from the *gypsy* transposon. **(B)** Bright and darkfield images of cuticles prepared from the different replacements: *gy, F7+gy, gy+F7, gy+F7 su(Hw)^-^*, CTCF^×4^, F7+CTCF^×4^, CTCF^×4^+F7 male flies. F7 replacement correspond to *wt.* (C) *Abd-B* expression in the CNS of stage 14 embryos. Staining with Engrailed to mark parasegment borders is shown in Fig. S1. All other designations are as in Fig. 1 and 2.

We generated two different combinations between *Fab-7* and *gy*: *F7+gy* and *gy+F7*. In the former, the *gypsy* insulator is next to the *iab-7* domain, while in the later the *gypsy* insulator is interposed between *Fab-7* and the *iab-6* domain. Fig. 4B shows that *F7+gy* combination for the most part supports bypass. With the exception of one or two bristles on the A6 sternite, the phenotype of *F7+gy* combination males is the same as *wt*. The trichomes on the A6 tergite are restricted to the anterior and ventral margins, while the sternite has the characteristic banana shape. Fab-7 also rescues the effects of gypsy in the embryonic CNS as *F7+gy* flies have expression pattern similar to *wt*. Consistent with the idea that order is important, the *gy+F7* combination does not support *iab-6* bypass and A6 resembles A5: the A6 sternite has multiple bristles and a quadrilateral shape while trichome hairs cover most of the A6 tergite. In addition, like *gy* alone, the trichomes hairs on the A5 tergite are densely packed as is typical of A4. Surprisingly, Abd-B expression in PS11 is suppressed in the *gy+F7* combination so that it resembles the pattern normally observed in PS10 (Fig. 4C, Fig. S1). In this instance, expression in the embryonic CNS parallels the alterations in segment identity observed in adult males.

The effects of *gy* on *Fab-7* bypass activity are expected to be due to the blocking activity of the *gy* insulator. To determine if this is the case, we introduced the *su(Hw)* mutation into the *gy+F7* background. Fig. 4B shows that inactivation of *su(Hw)* fully rescues the bypass defects of *gy+F7* in the adult male cuticle.

### Multimerized CTCF binding sites can also disrupt *Fab-7* bypass activity

These results argue that heterologous insulators can disrupt bypass activity when placed between the *iab-6* domain and *Fab-7*. However, if the heterologous insulator is adjacent to the *iab-7* regulatory domain, bypass activity is retained. To confirm this conclusion, we tested a completely artificial boundary, CTCF^×4^, which consist of four dCTCF binding sites. CTCF^×4^ alone blocks crosstalk between *iab-6* and *iab-7*, but does not allow the *iab-6* domain to properly activate *Abd-B* in PS11/A6 [48]. The A6 tergite is almost completely covered in trichome hairs, while the sternite is somewhat misshapen and covered in bristles. Like the *gy* replacement, it also interferes with the functioning of the *iab-5* domain in A5. The trichome hairs on the A5 tergite are densely packed like A4 and there are small patches of cuticle that lack pigmentation. The bypass defects seen for CTCF^×4^ in both A6 and A5 are rescued by F7+CTCF^×4^ combination: the morphology of A6 resembles *wt* (Fig. 4B). In contrast, F7 bypass activity is lost when CTCF^×4^ is placed between *iab-6* and the F7. In this case the morphology of A6 resembles that seen with CTCF^×4^ alone.

In the embryonic CNS, Abd-B expression in both PS10 and PS11 is reduced by CTCF^×4^ alone (Fig. 4C, Fig. S1). For CTCF^×4^+F7 combination we observed a complete loss of Abd-B expression in both PS10 and PS11. Addition of *Fab-7* before CTCF^×4^(F7+CTCF^×4^) restored Abd-B expression in PS10. Whereas Abd-B expression in PS11 was still reduced compared to *wt* (Fig. 4C).

### The bypass function of *Fab-7* dHS1 is blocked by multimerized Pita sites

As described in the Introduction, we found that the bypass defects of *Pita^×5^* can be rescued when it is combine with the 242 bp *Fab-7* dHS1 fragment (Fig. 1A, 5A) [49]. In these experiments dHS1 was placed next to the *iab-6* domain, while *Pita^×5^* bordered the *iab-7* domain (Fig. 5A). According to the results in the previous sections, the bypass function of dHS1 should be disrupted when the order is reversed and *Pita^×5^* is interposed between the *iab-6* regulatory domain and dHS1. Fig. 5B shows that this prediction holds. In the *Pita^×5^* replacement the A6 tergite is nearly covered in trichome hairs, while the sternite is misshapen and unlike *wt* has bristles. As was observed for both the *gy* and CTCF^×4^ replacements, there is also evidence of a LOF transformation of PS10/A5 towards PS9/A4. While these LOF phenotypes are rescued in the *dHS1+Pita^×5^*replacement and both A6 and A5 resemble *wt*, this is not true for the *Pita^×5^+dHS1* combination: trichomes cover most of the A6 tergite while the trichomes in the A5 tergite are densely packed like A4. Though the *Pita^×5^+dHS1* fails to rescue the LOF transformations of the A5 and A6 tergites, the A6 sternite has a banana-like shape but is covered in bristles. The bypass defects in *Pita^×5^+dHS1* are not rescued by introducing a second Pita multimer (*Pita^×5^+dHS1+Pita^×5^*) in between *dHS1* and the *iab-7* regulatory domain. The phenotypes of the adult cuticle correlate with the pattern of expression of *Abd-B* in the embryonic CNS. Fig. 5C (Fig. S1) shows that *Abd-B* expression in PS11 and PS10 in *dHS1+Pita^×5^* is similar to that of *wt*, while both the *Pita^×5^+dHS1* and *Pita^×5^+dHS1+Pita^×5^* combinations resemble *Pita^×5^* in that there is only little Abd-B in PS11 while *Abd-B* expression appears to be absent in PS10.

**Fig. 5.**
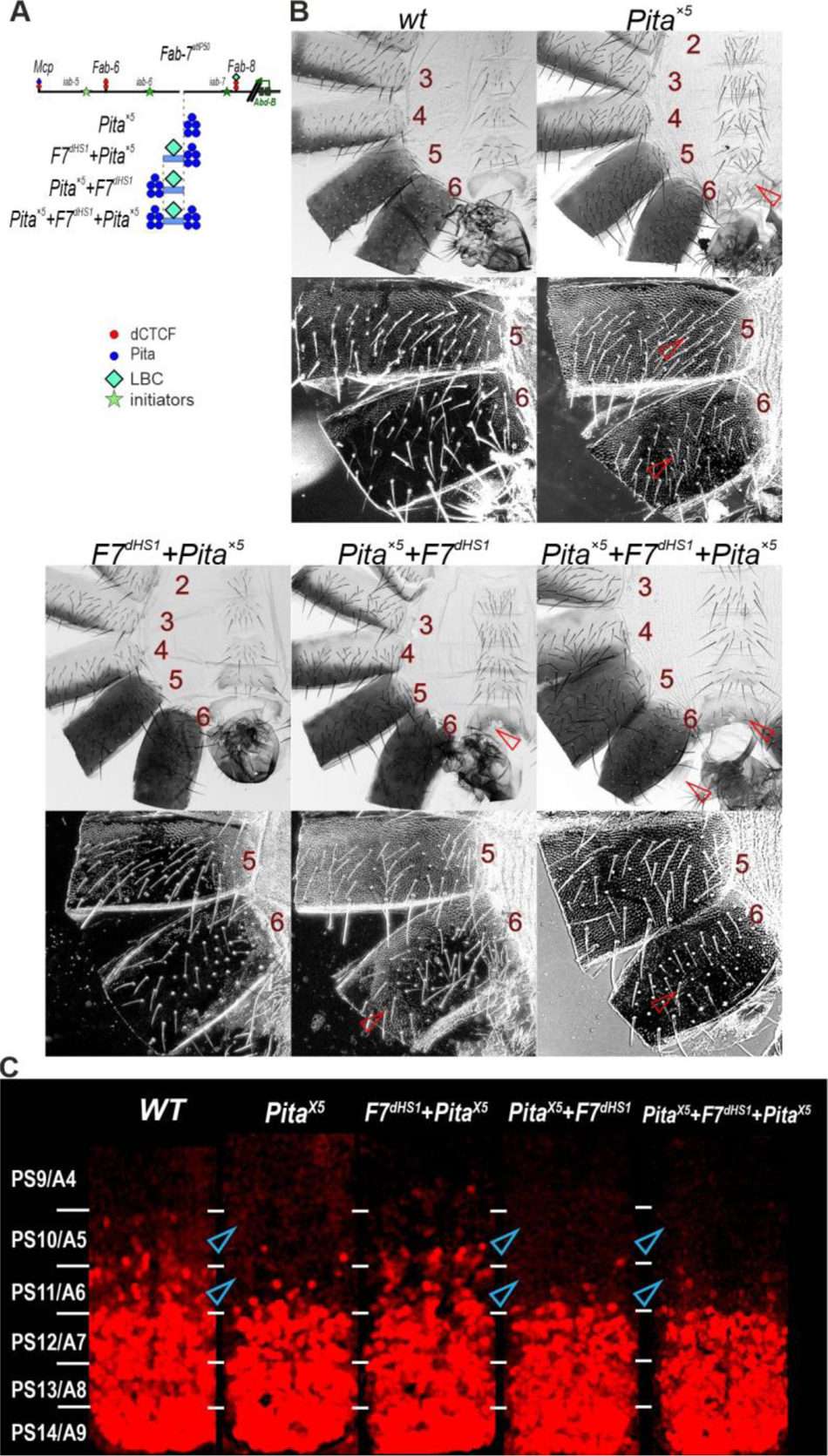
Multimerized Pita sites disrupt the bypass activity of *Fab-7* dHS1. **(A)** Schematic presentation of *Fab-7* substitutions with different combinations of the *Pita^×5^*and *F7^dHS1^*. **(B)** Images of cuticles prepared from *Pita^×5^, F7^dHS1^+Pita^×5^* (phenotype similar to *wt*), *Pita^×5^+F7^dHS1^*, and *Pita^×5^+F7^dHS1^+Pita^×5^* male flies. **(C)** *Abd-B* expression in the CNS of stage 14 embryos. **(D)** Images of cuticles prepared from *wt,*, *Pita^×5^, F7^dHS1^+Pita^×5^*(phenotype similar to *wt*), *Pita^×5^+F7^dHS1^*, and *Pita^×5^+F7^dHS1^+Pita^×5^* male flies.. All designations are as in Fig. 1 and 2.

### Multimerized Pita sites disrupt *Fab-8* bypass activity

To confirm that the position dependent effects of *Pita^×5^* on bypass activity are not restricted to the *Fab-7* dHS1, we generated two different *Pita^×5^* combinations with *Fab-8*, *F8+Pita^×5^* and *Pita^×5^+F8* (Fig. 6A). In the former the *Fab-8* bypass element is adjacent to *iab-6*, while in the latter *Pita^×5^* is interposed between *iab-6* and *Fab-8*. As was observed for the *dHS1+Pita^×5^*combination, the morphology of A6 in *F8+Pita^×5^* males resembles *wt,* excluding a few bristles on sternite (Fig. 6B). However, when *Pita^×5^*is placed between *iab-6* and F8, bypass activity is lost and the morphology of A6 is the same as that of A5. The effects on cuticular differentiation seen in the adult are reflected in the pattern of *Abd-B* expression in the embryonic CNS. As shown in Fig. 6C (Fig. S1), Abd-B expression in PS11 and PS10 in F8 and *F8+Pita^×5^* is the same as *wt*. In contrast, in *Pita^×5^+F8* and *Pita^×5^ Abd-B* expression in PS11 is substantially reduced and appears to be absent in PS10.

**Fig. 6.**
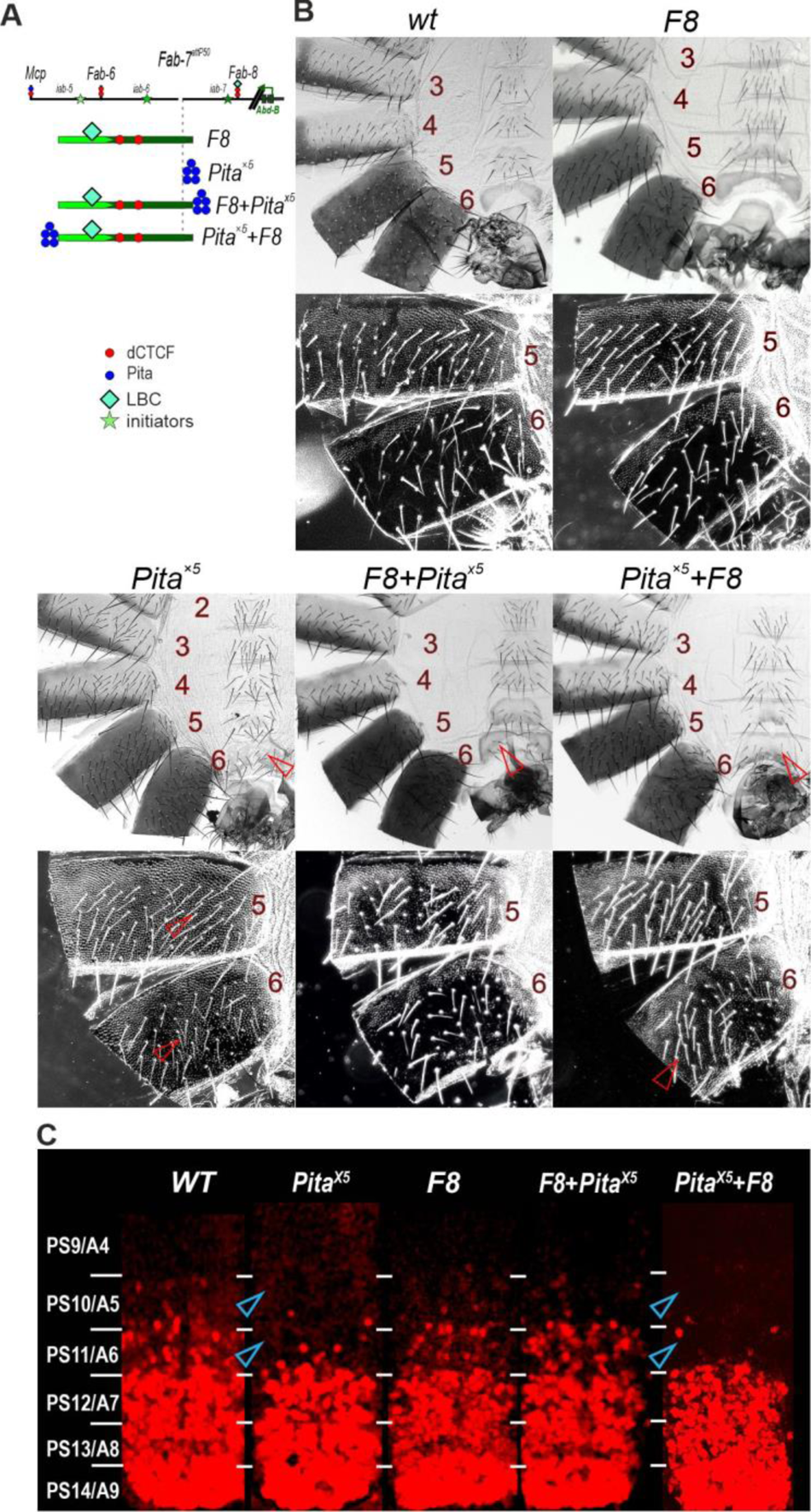
Multimerized Pita sites disrupt *Fab-8* bypass activity. **A)** Schematic presentation of *Fab-7* substitutions. **B)** Images of cuticles prepared from *wt, F8* (phenotype similar to *wt*), *F8+Pita^×5^,* and *Pita^×5^+ F8* male flies. All designations are as in Fig. 2 and 3. **(C)** *Abd-B* expression in the CNS of stage 14 embryos. **(D)** Images of cuticles prepared from *wt, F8* (phenotype similar to *wt*), *F8+Pita^×5^,* and *Pita^×5^+ F8* male flies. All designations are as in Fig. 1 and 2.

## DISCUSSION

Distant interactions between enhancers/silencers and their regulatory targets are a common feature of gene regulation in complex multicellular organisms [21,62–65]. These interactions can occur over distances of kbs to Mbs and can span one or more intervening TADs together with their associated genes and boundaries. As boundaries normally restrict interactions to regulatory elements and genes in the same domain, the blocking activity of intervening boundaries must be bypassed. However, the bypass mechanism must also ensure specificity so that the looping enhancers/silencers do not impact the functioning of intervening TADs. One well studied context for analyzing long distance regulatory interactions that must circumvent intervening boundary elements is the *Abd-B* region of the *Drosophila* BX-C. Three of the *Abd-B* regulatory domains, *iab-5*, *iab-6* and *iab-7* are separated from the promoter *Abd-B* by three (*Fab-6*, *Fab-7*, *Fab-8*), two (*Fab-7, Fab-8*) or one (*Fab-8*) boundary element, respectively. For this reason, the *Abd-B* boundaries have two functions – they block crosstalk between neighboring regulatory domains, and at the same time actively facilitate long distance communication between the regulatory domains and their target, *Abd-B* [65]. This model is supported by the interaction between the *Fab-7* or *Fab-8* boundaries and the *Abd-B* promoter region found in MicroC studies of embryos [66].

In previous *Fab-7* replacement studies, we found that the orientation of the *Fab-8* boundary affects interactions between *iab-6* and *Abd-B* promoter. Communication is observed when the *Fab-8* replacement is in the same orientation as the endogenous *Fab-8* boundary; however, if the orientation of *Fab-8* is reversed only blocking activity is observed [48]. Since we observed a similar orientation dependence between *Fab-8* and the AB-I promoter element in transgene insulator bypass assays [59], we thought that a similar mechanism was at play in the *Fab-7* replacement assay. We subsequently discovered that *Fab-8* consists of two elements: a 165 bp element, F8^165^, whose primary function is bypass, and a 209 bp element, F8^209^, that functions as an insulator [54]. Here we show that the orientation of the F8^165^ and F8^209^ elements is not important; instead, their relative order matters. This requirement is not due to some special property of the F8^165^ bypass element as similar results are obtained for composite boundaries containing *Fab-7* and different insulators (*F8^209^, gy* or CTCF^×4^). When *Fab-7* flanks the *iab-6* domain (*F7+gy* or F7+CTCF^×4^) the composite boundary has both blocking and bypass activity. However, when the insulator elements are between the *iab-6* domain and *Fab-7* bypass activity is lost. We also tested *Fab-7* dHS1 which has bypass activity when combined with multimerized sites for the architectural protein Pita. As was observed for the *Fab-8* bypass element, dHS1 must be next to the *iab-6* domain to mediate bypass.

It is interesting to note that the effects of elements that have insulating activity on *Abd-B* regulation are not identical. The *Fab-8* insulating element, F8^209^, induces a LOF transformation of PS11/A6 towards PS10/A5. However, in adult males the transformation is incomplete as both the tergite and the sternite retain A6-like features. In contrast, the CTCF^×4^, *Pita^×5^* and *gy* replacements induce a more complete transformation of A6 into A5. In all three replacements the A6 tergite is mostly covered in trichomes. The sternite in CTCF^×4.^ is misshapen and has bristles, while for both *Pita^×5^* and *gy* the sternite is similar to A5. Moreover, all three of these boundaries also impact the development of the A5 segment. The trichomes on the A5 tergite are densely packed like those in the A4 tergite, and there are patches of unpigmented cuticle. These finding indicate that unlike F8^209^, these boundaries are able to interfere with *iab-5* regulation of *Abd-B*. On the other hand, the bypass elements we tested were able to circumvent their blocking activity and restore normal morphology to both A5 and A6.

In previous studies, we tested the ability of dHS1 to rescue the bypass defects of multimerized CTCF^×4^ and *Su(Hw)^×4^* sites [49]. Like multimerized *Pita^×5^* sites, both not only block *iab-6* from regulating *Abd-B* in PS11/A6, but they also interfere with *iab-5* regulation of *Abd-B* in PS10/A5. Also, like multimerized *Pita^×5^* sites, these bypass defects can be rescued by the addition of dHS1. These findings would suggest that the bypass activity *Fab-7* dHS1 (and likely also F8^165^) does not depend upon some special property of the insulator element that makes it permissive to bypass.

While the studies reported here together with our previous work [49,50,54] indicate that the bypass elements in the *Fab-7* and *Fab-8* boundaries enable regulatory domains to “jump over” intervening boundaries and activate *Abd-B*, it remains unknown when and where these interactions take place. On the one hand, such interactions could be established as boundaries are assembled and TADs are formed during the early nuclear division cycles and then remain in place through the rest of development. On the other hand, these interactions might be coordinated with the activity state of the regulatory domain (Fig. 7A). In PS10/A5 and in more anterior parasegments/segments, the *Fab-7* bypass element, dHS1, might be inactive and not mediate contacts between *iab-6* and *Abd-B*. In contrast, in PS11/A6, where the *iab-6* regulatory domain is active, the *Fab-7* bypass element would be activated and function to bring the *iab-6* enhancers into contact with the *Abd-B* gene. A connection between the activity state of the *iab-6* domain and bypass activity could explain why the bypass elements needs to be next to the domain that is driving *Abd-B* expression and why it is unable to function when a boundary element is interposed between it and the active *iab-6* domain (Fig. 7B,C). With respect to *Abd-B* regulation, the idea that bypass activity is subject to regulation would seem to make sense. For one it would mean that distal active regulatory domains would not need to compete with more proximal PcG silenced domains for interactions with the *Abd-B* promoters. Additionally, domains that are off and subject to Polycomb silencing would not be brought into close proximity to the *Abd-B* promoter region where they could potentially inhibit expression.

**Fig. 7.**
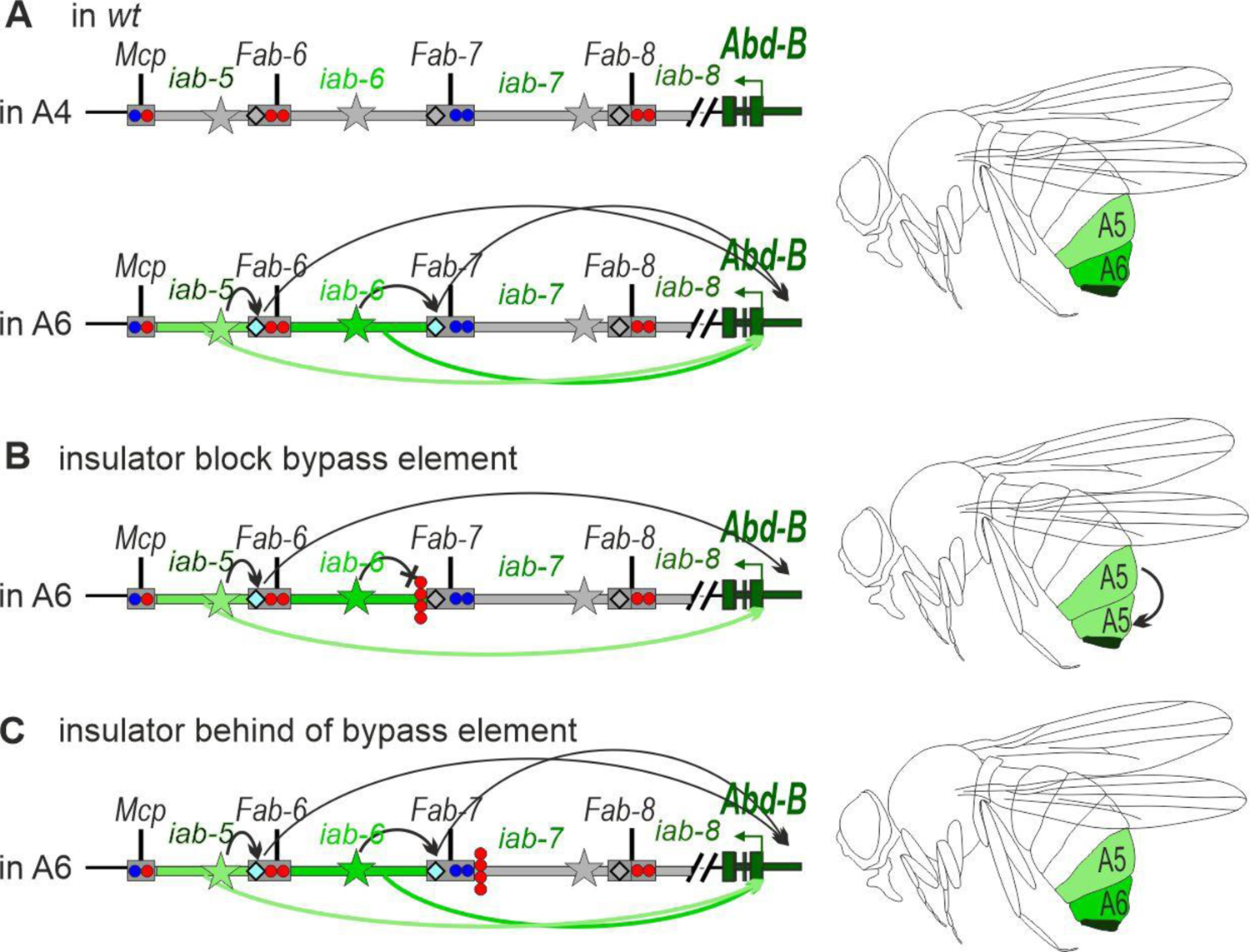
Model of the functional role of the *iab-6* activation in the regulation of distance interaction between the *Fab-7* boundary and the *Abd-B* promoter region. (A) In *wt* in A4 segment *Abd-B* regulatory region is inactive, initiators are repressed and the *Fab-*boundaries does not interact with the *Abd-B* promoter region. Activation of the *iab-6* domain in PS11/A6 results in the stimulation of the *Fab-7* bypass module. The interaction between the *Fab-7* boundary and *Abd-B* promoter region facilitates enhancer-promoter communication. **(B)** An insulator inserted between the *iab-6* enhancers and the bypass element (proximal side) blocks interaction with the *Abd-B* promoter region. **(C)** The insulator inserted on the distal side of the *Fab-7* boundary does not interfere with correct stimulation of the bypass element.

An important question is whether there are other contexts which utilize “bypass” elements like those in the *Abd-B* region of BX-C to mediate long distance enhancer:promoter interactions. Eagen et al., 2017 [67] found that in developmental loci like *engrailed*-*invected* and the Antennapedia complex that contain two or more PREs, the PREs interact with each other, forming interaction dots in HiC experiments. More recent studies [66,68] showed that PREs in these loci function as enhancer-promoter “tethering elements.” By pairing with each other, they help to physically link distant enhancers to their target genes. For example, in the *knirps* locus, there are PREs located just upstream of the *knirps* (*krp* and *knirps-like* (*knpl*) genes (see double headed arrow in Fig. S2). The PREs interact with each other and bring the two genes together in 3D space (see double headed arrow in Fig. S2). Deletion of the PRE upstream of the *knrl* gene eliminates the long-distance PRE-PRE contacts and also disrupts expression of the *knr* gene located some 70 kb away.

Interestingly, the ChIP signatures (Fig. S2) of the tethering elements identified in [66,68] share features with those found in the *Fab-7* bypass element, dHS1, and the *Fab-8* bypass element F8^165^ (Fig. S3). Like the *Fab-7* and *Fab-8* bypass elements, GAF and CLAMP are found in most of the tethering elements. In the case of dHS1, we showed that its bypass activity is sensitive to reductions dose of both GAF (*Trl*) and CLAMP [49,54]. Since these ChIP signatures are found at sequences that are known to be bound by the LBC in embryonic nuclear extracts, it seems likely that the tethering elements identified by Levo et al., (2022) and Batut et al., (2022) [66,68] will also interact with the LBC in nuclear extracts. In this case both inter- and intra-TAD enhancer-promoter interactions maybe mediated by the LBC.

Boundary pairing interactions in flies are typically orientation dependent and this means that TADs can be either stem-loops or circle-loops [58]. The topology of the loop that is generated by boundary:boundary pairing is important as it can determine which elements (enhancers/silencers/promoters) are in close proximity to each other [15,58,69]. In this respect it is interesting that the bypass and blocking elements in the *Fab-8* boundary can be reversed without disrupting bypass. One would have expected that inverting the two elements (but keeping the order the same) would switch the loop topology from the predicted circle-loop to a stem-loop and this would disrupt bypass activity as is observed in transgene assays [15,58,69]. Since bypass is still observed, this could mean that loop topology is unaltered. Alternatively, the bypass element might be able to mediate enhancer/silencer: promoter interactions independent of pairing orientation. In this respect it is interesting that the *Fab-7* boundary, whose activity depends upon two LBC binding sequences, dHS1 and HS3, shows limited orientation dependence. Further studies will be required to investigate this problem.

## MATERIALS AND METHODS

### Generation of transgenic lines carrying different deletions and insertions

The strategy of the *Fab-7* replacement lines is described in detail in [48,55]. To introduce *gy+F7* into the *su(Hw*)^-^ null background, we combined *su(Hw)v/su(Hw)^e04061^* and *gy+F7* as described previously [70]. To introduce *F8^209^+F7 into the dctcf* null background, we combined the GE24185 mutation and F8^209^+F7 as described previously [60,61].

### Antibody staining in embryos

Embryos were stained following standard protocols. Embryos were collected for 19 hours. Primary antibodies were mouse monoclonal anti-Abd-B at 1:40 dilution (1A2E9, generated by S.Celniker, deposited to the Developmental Studies Hybridoma Bank) and polyclonal rabbit anti-Engrailed at 1:500 dilution (kindly provided to us by Judith Kassis). Secondary antibodies were goat anti-mouse Alexa Fluor 546 and anti-rabbit Alexa Fluor 488 (Thermo Fisher Scientific) at 1:500 dilution. Stained embryos were mounted in Vectashield. At least 40 stage 15 embryos of each genotype were examined. The most representative embryo for each transgenic line was selected for presentation in the manuscript. Images were acquired on Leica Stellaris 5 confocal microscope and processed using ImageJ 1.50c4

### Cuticle preparations

Cuticle preparations were carried out as described in [50]. Phenotypes depicted are representative of the genotypes shown. For each transgenic line, visual analysis was performed on approximately 50 males. Sometimes occasional variances are observed in the exact number of bristles and the exact pattern of trichomes. The most phenotypically different 3-4 males were selected for preparation of photos with the abdominal cuticles. If there were no statistically significant differences in cuticles, we attempted to select an average representative cuticle for display.

## ACKNOWLEDGEMENTS

We thank Farhod Hasanov for fly injections, Kate O’Connor-Giles for the pHD-DsRed plasmid (Addgene plasmid #51434) and pAc5-Cas9 containing plasmid (Addgene plasmid # 62209), Bloomington Drosophila Stock Center for Drosophila lines.

## Funding

This work (all functional and morphological analysis) was supported by the Russian Science Foundation (19-14-00103 to O.K.). Part of this work (genome editing procedure) was supported by grant 075-15-2019-1661 from the Ministry of Science and Higher Education of the Russian Federation. P.S. acknowledges support from the National Institutes of Health (R35 GM126975).

## Author contributions

Conceptualization: P.G., O.K.; Methodology: A.I., N.P., O.K.; Validation: P.G., O.K.; Formal analysis: P.S., P.G., O.K.; Investigation: A.I., N.P., O.K.; Resources: O.K.; Data curation: P.G., O.K.; Writing - original draft: P.S., P.G., O.K.; Writing - review & editing: P.S., P.G., O.K.; Visualization: A.I., N.P., O.K.; Supervision: P.G., O.K., P.S.; Project administration: P.G., O.K.; Funding acquisition: O.K., P.S.

## Competing interests

The authors declare no competing or financial interests.

## Data and materials availability

All data needed to evaluate the conclusions in the paper are present in the paper and/or the Supplementary Materials. Additional data related to this paper may be requested from the authors.

## Notes

### Competing Interest Statement

The authors have declared no competing interest.

## REFERENCES

1. Cavalheiro GR, Pollex T, Furlong EE. 2021 To loop or not to loop: what is the role of TADs in enhancer function and gene regulation? Curr Opin Genet Dev 67, 119–129. (doi:10.1016/j.gde.2020.12.015)

2. Hafner A, Boettiger A. 2022 The spatial organization of transcriptional control. Nat Rev Genet (doi:10.1038/s41576-022-00526-0)

3. Kyrchanova O, Georgiev P. 2021 Mechanisms of Enhancer-Promoter Interactions in Higher Eukaryotes. Int J Mol Sci 22, E671. (doi:10.3390/ijms22020671)

4. Peterson SC, Samuelson KB, Hanlon SL. 2021 Multi-Scale Organization of the Drosophila melanogaster Genome. Genes (Basel) 12, 817. (doi:10.3390/genes12060817)

5. Shukla V, Cetnarowska A, Hyldahl M, Mandrup S. 2022 Interplay between regulatory elements and chromatin topology in cellular lineage determination. Trends Genet 38, 1048–1061. (doi:10.1016/j.tig.2022.05.011)

6. van Steensel B, Furlong EEM. 2019 The role of transcription in shaping the spatial organization of the genome. Nat Rev Mol Cell Biol 20, 327–337. (doi:10.1038/s41580-019-0114-6)

7. Rao SSP et al. 2014 A 3D map of the human genome at kilobase resolution reveals principles of chromatin looping. Cell 159, 1665–1680. (doi:10.1016/j.cell.2014.11.021)

8. Bag I, Chen S, Rosin LF, Chen Y, Liu C-Y, Yu G-Y, Lei EP. 2021 M1BP cooperates with CP190 to activate transcription at TAD borders and promote chromatin insulator activity. Nat Commun 12, 4170. (doi:10.1038/s41467-021-24407-y)

9. Ramírez F, Bhardwaj V, Arrigoni L, Lam KC, Grüning BA, Villaveces J, Habermann B, Akhtar A, Manke T. 2018 High-resolution TADs reveal DNA sequences underlying genome organization in flies. Nat Commun 9, 189. (doi:10.1038/s41467-017-02525-w)

10. Wang Q, Sun Q, Czajkowsky DM, Shao Z. 2018 Sub-kb Hi-C in D. melanogaster reveals conserved characteristics of TADs between insect and mammalian cells. Nat Commun 9, 188. (doi:10.1038/s41467-017-02526-9)

11. Zolotarev N et al. 2016 Architectural proteins Pita, Zw5,and ZIPIC contain homodimerization domain and support specific long-range interactions in Drosophila. Nucleic Acids Res 44, 7228–7241. (doi:10.1093/nar/gkw371)

12. Kyrchanova OV, Bylino OV, Georgiev PG. 2022 Mechanisms of enhancer-promoter communication and chromosomal architecture in mammals and Drosophila. Front Genet 13, 1081088. (doi:10.3389/fgene.2022.1081088)

13. Barges S, Mihaly J, Galloni M, Hagstrom K, Müller M, Shanower G, Schedl P, Gyurkovics H, Karch F. 2000 The Fab-8 boundary defines the distal limit of the bithorax complex iab-7 domain and insulates iab-7 from initiation elements and a PRE in the adjacent iab-8 domain. Development 127, 779–790.

14. Hagstrom K, Muller M, Schedl P. 1996 Fab-7 functions as a chromatin domain boundary to ensure proper segment specification by the Drosophila bithorax complex. Genes Dev 10, 3202–3215. (doi:10.1101/gad.10.24.3202)

15. Kyrchanova O, Chetverina D, Maksimenko O, Kullyev A, Georgiev P. 2008 Orientation-dependent interaction between Drosophila insulators is a property of this class of regulatory elements. Nucleic Acids Res 36, 7019–7028. (doi:10.1093/nar/gkn781)

16. Kyrchanova O, Maksimenko O, Stakhov V, Ivlieva T, Parshikov A, Studitsky VM, Georgiev P. 2013 Effective blocking of the white enhancer requires cooperation between two main mechanisms suggested for the insulator function. PLoS Genet 9, e1003606. (doi:10.1371/journal.pgen.1003606)

17. Pérez-Lluch S, Cuartero S, Azorín F, Espinàs ML. 2008 Characterization of new regulatory elements within the Drosophila bithorax complex. Nucleic Acids Res 36, 6926–6933. (doi:10.1093/nar/gkn818)

18. Savitsky M, Kim M, Kravchuk O, Schwartz YB. 2016 Distinct Roles of Chromatin Insulator Proteins in Control of the Drosophila Bithorax Complex. Genetics 202, 601–617. (doi:10.1534/genetics.115.179309)

19. Zhou J, Barolo S, Szymanski P, Levine M. 1996 The Fab-7 element of the bithorax complex attenuates enhancer-promoter interactions in the Drosophila embryo. Genes Dev 10, 3195–3201. (doi:10.1101/gad.10.24.3195)

20. Galouzis CC, Furlong EEM. 2022 Regulating specificity in enhancer-promoter communication. Curr Opin Cell Biol 75, 102065. (doi:10.1016/j.ceb.2022.01.010)

21. Jerković I, Szabo Q, Bantignies F, Cavalli G. 2020 Higher-Order Chromosomal Structures Mediate Genome Function. J Mol Biol 432, 676–681. (doi:10.1016/j.jmb.2019.10.014)

22. Amândio AR, Beccari L, Lopez-Delisle L, Mascrez B, Zakany J, Gitto S, Duboule D. 2021 Sequential in cis mutagenesis in vivo reveals various functions for CTCF sites at the mouse HoxD cluster. Genes Dev 35, 1490–1509. (doi:10.1101/gad.348934.121)

23. Kane L, Williamson I, Flyamer IM, Kumar Y, Hill RE, Lettice LA, Bickmore WA. 2022 Cohesin is required for long-range enhancer action at the Shh locus. Nat Struct Mol Biol 29, 891–897. (doi:10.1038/s41594-022-00821-8)

24. Symmons O, Pan L, Remeseiro S, Aktas T, Klein F, Huber W, Spitz F. 2016 The Shh Topological Domain Facilitates the Action of Remote Enhancers by Reducing the Effects of Genomic Distances. Dev Cell 39, 529–543. (doi:10.1016/j.devcel.2016.10.015)

25. Lewis EB. 1978 A gene complex controlling segmentation in Drosophila. Nature 276, 565–570. (doi:10.1038/276565a0)

26. Duncan I. 1987 The bithorax complex. Annu Rev Genet 21, 285–319. (doi:10.1146/annurev.ge.21.120187.001441)

27. Peifer M, Karch F, Bender W. 1987 The bithorax complex: control of segmental identity. Genes Dev 1, 891–898. (doi:10.1101/gad.1.9.891)

28. Maeda RK, Karch F. 2015 The open for business model of the bithorax complex in Drosophila. Chromosoma 124, 293–307. (doi:10.1007/s00412-015-0522-0)

29. Kyrchanova O, Mogila V, Wolle D, Magbanua JP, White R, Georgiev P, Schedl P. 2015 The boundary paradox in the Bithorax complex. Mech Dev 138 **Pt** **2**, 122–132. (doi:10.1016/j.mod.2015.07.002)

30. Bender WW. 2020 Molecular Lessons from the Drosophila Bithorax Complex. Genetics 216, 613– 617. (doi:10.1534/genetics.120.303708)

31. Mihaly J, Barges S, Sipos L, Maeda R, Cléard F, Hogga I, Bender W, Gyurkovics H, Karch F. 2006 Dissecting the regulatory landscape of the Abd-B gene of the bithorax complex. Development 133, 2983–2993. (doi:10.1242/dev.02451)

32. Ho MCW, Schiller BJ, Goetz SE, Drewell RA. 2009 Non-genic transcription at the Drosophila bithorax complex functional activity of the dark matter of the genome. Int J Dev Biol 53, 459–468. (doi:10.1387/ijdb.082647mh)

33. Starr MO, Ho MCW, Gunther EJM, Tu Y-K, Shur AS, Goetz SE, Borok MJ, Kang V, Drewell RA. 2011 Molecular dissection of cis-regulatory modules at the Drosophila bithorax complex reveals critical transcription factor signature motifs. Dev Biol 359, 290–302. (doi:10.1016/j.ydbio.2011.07.028)

34. Shimell MJ, Simon J, Bender W, O’Connor MB. 1994 Enhancer point mutation results in a homeotic transformation in Drosophila. Science 264, 968–971. (doi:10.1126/science.7909957)

35. Iampietro C, Gummalla M, Mutero A, Karch F, Maeda RK. 2010 Initiator elements function to determine the activity state of BX-C enhancers. PLoS Genet 6, e1001260. (doi:10.1371/journal.pgen.1001260)

36. Drewell RA, Nevarez MJ, Kurata JS, Winkler LN, Li L, Dresch JM. 2014 Deciphering the combinatorial architecture of a Drosophila homeotic gene enhancer. Mech Dev 131, 68–77. (doi:10.1016/j.mod.2013.10.002)

37. Casares F, Sánchez-Herrero E. 1995 Regulation of the infraabdominal regions of the bithorax complex of Drosophila by gap genes. Development 121, 1855–1866.

38. Postika N, Schedl P, Georgiev P, Kyrchanova O. 2021 Redundant enhancers in the iab-5 domain cooperatively activate Abd-B in the A5 and A6 abdominal segments of Drosophila. Development 148, dev199827. (doi:10.1242/dev.199827)

39. Kennison JA. 1993 Transcriptional activation of Drosophila homeotic genes from distant regulatory elements. Trends Genet 9, 75–79. (doi:10.1016/0168-9525(93)90227-9)

40. Müller J, Kassis JA. 2006 Polycomb response elements and targeting of Polycomb group proteins in Drosophila. Curr Opin Genet Dev 16, 476–484. (doi:10.1016/j.gde.2006.08.005)

41. Simon JA, Kingston RE. 2009 Mechanisms of polycomb gene silencing: knowns and unknowns. Nat Rev Mol Cell Biol 10, 697–708. (doi:10.1038/nrm2763)

42. Galloni M, Gyurkovics H, Schedl P, Karch F. 1993 The bluetail transposon: evidence for independent cis-regulatory domains and domain boundaries in the bithorax complex. EMBO J 12, 1087–1097.

43. Karch F, Galloni M, Sipos L, Gausz J, Gyurkovics H, Schedl P. 1994 Mcp and Fab-7: molecular analysis of putative boundaries of cis-regulatory domains in the bithorax complex of Drosophila melanogaster. Nucleic Acids Res 22, 3138–3146. (doi:10.1093/nar/22.15.3138)

44. Bender W, Lucas M. 2013 The border between the ultrabithorax and abdominal-A regulatory domains in the Drosophila bithorax complex. Genetics 193, 1135–1147. (doi:10.1534/genetics.112.146340)

45. Gyurkovics H, Gausz J, Kummer J, Karch F. 1990 A new homeotic mutation in the Drosophila bithorax complex removes a boundary separating two domains of regulation. EMBO J 9, 2579–2585.

46. Bowman SK, Deaton AM, Domingues H, Wang PI, Sadreyev RI, Kingston RE, Bender W. 2014 H3K27 modifications define segmental regulatory domains in the Drosophila bithorax complex. Elife 3, e02833. (doi:10.7554/eLife.02833)

47. Postika N, Schedl P, Georgiev P, Kyrchanova O. 2021 Mapping of functional elements of the Fab-6 boundary involved in the regulation of the Abd-B hox gene in Drosophila melanogaster. Sci Rep 11, 4156. (doi:10.1038/s41598-021-83734-8)

48. Kyrchanova O, Mogila V, Wolle D, Deshpande G, Parshikov A, Cléard F, Karch F, Schedl P, Georgiev P. 2016 Functional Dissection of the Blocking and Bypass Activities of the Fab-8 Boundary in the Drosophila Bithorax Complex. PLoS Genet 12, e1006188. (doi:10.1371/journal.pgen.1006188)

49. Kyrchanova O, Sabirov M, Mogila V, Kurbidaeva A, Postika N, Maksimenko O, Schedl P, Georgiev P. 2019 Complete reconstitution of bypass and blocking functions in a minimal artificial Fab-7 insulator from Drosophila bithorax complex. Proc Natl Acad Sci U S A 116, 13462–13467. (doi:10.1073/pnas.1907190116)

50. Postika N, Metzler M, Affolter M, Müller M, Schedl P, Georgiev P, Kyrchanova O. 2018 Boundaries mediate long-distance interactions between enhancers and promoters in the Drosophila Bithorax complex. PLoS Genet 14, e1007702. (doi:10.1371/journal.pgen.1007702)

51. Hogga I, Mihaly J, Barges S, Karch F. 2001 Replacement of Fab-7 by the gypsy or scs insulator disrupts long-distance regulatory interactions in the Abd-B gene of the bithorax complex. Mol Cell 8, 1145– 1151. (doi:10.1016/s1097-2765(01)00377-x)

52. Kyrchanova O, Zolotarev N, Mogila V, Maksimenko O, Schedl P, Georgiev P. 2017 Architectural protein Pita cooperates with dCTCF in organization of functional boundaries in Bithorax complex. Development 144, 2663–2672. (doi:10.1242/dev.149815)

53. Kyrchanova O et al. 2018 The bithorax complex iab-7 Polycomb response element has a novel role in the functioning of the Fab-7 chromatin boundary. PLoS Genet 14, e1007442. (doi:10.1371/journal.pgen.1007442)

54. Kyrchanova O, Wolle D, Sabirov M, Kurbidaeva A, Aoki T, Maksimenko O, Kyrchanova M, Georgiev P, Schedl P. 2019 Distinct Elements Confer the Blocking and Bypass Functions of the Bithorax Fab-8 Boundary. Genetics 213, 865–876. (doi:10.1534/genetics.119.302694)

55. Wolle D, Cleard F, Aoki T, Deshpande G, Schedl P, Karch F. 2015 Functional Requirements for Fab-7 Boundary Activity in the Bithorax Complex. Mol Cell Biol 35, 3739–3752. (doi:10.1128/MCB.00456-15)

56. Hagstrom K, Muller M, Schedl P. 1997 A Polycomb and GAGA dependent silencer adjoins the Fab-7 boundary in the Drosophila bithorax complex. Genetics 146, 1365–1380. (doi:10.1093/genetics/146.4.1365)

57. Mishra RK, Mihaly J, Barges S, Spierer A, Karch F, Hagstrom K, Schweinsberg SE, Schedl P. 2001 The iab-7 polycomb response element maps to a nucleosome-free region of chromatin and requires both GAGA and pleiohomeotic for silencing activity. Mol Cell Biol 21, 1311–1318. (doi:10.1128/MCB.21.4.1311-1318.2001)

58. Chetverina D, Fujioka M, Erokhin M, Georgiev P, Jaynes JB, Schedl P. 2017 Boundaries of loop domains (insulators): Determinants of chromosome form and function in multicellular eukaryotes. Bioessays 39. (doi:10.1002/bies.201600233)

59. Kyrchanova O, Toshchakov S, Podstreshnaya Y, Parshikov A, Georgiev P. 2008 Functional interaction between the Fab-7 and Fab-8 boundaries and the upstream promoter region in the Drosophila Abd-B gene. Mol Cell Biol 28, 4188–4195. (doi:10.1128/MCB.00229-08)

60. Kyrchanova O, Klimenko N, Postika N, Bonchuk A, Zolotarev N, Maksimenko O, Georgiev P. 2021 Drosophila architectural protein CTCF is not essential for fly survival and is able to function independently of CP190. Biochim Biophys Acta Gene Regul Mech 1864, 194733. (doi:10.1016/j.bbagrm.2021.194733)

61. Mohan M et al. 2007 The Drosophila insulator proteins CTCF and CP190 link enhancer blocking to body patterning. EMBO J 26, 4203–4214. (doi:10.1038/sj.emboj.7601851)

62. Ray-Jones H, Spivakov M. 2021 Transcriptional enhancers and their communication with gene promoters. Cell Mol Life Sci 78, 6453–6485. (doi:10.1007/s00018-021-03903-w)

63. Panigrahi A, O’Malley BW. 2021 Mechanisms of enhancer action: the known and the unknown. Genome Biol 22, 108. (doi:10.1186/s13059-021-02322-1)

64. Jindal GA, Farley EK. 2021 Enhancer grammar in development, evolution, and disease: dependencies and interplay. Dev Cell 56, 575–587. (doi:10.1016/j.devcel.2021.02.016)

65. Kyrchanova O, Sokolov V, Georgiev P. 2023 Mechanisms of Interaction between Enhancers and Promoters in Three Drosophila Model Systems. Int J Mol Sci 24, 2855. (doi:10.3390/ijms24032855)

66. Batut PJ, Bing XY, Sisco Z, Raimundo J, Levo M, Levine MS. 2022 Genome organization controls transcriptional dynamics during development. Science 375, 566–570. (doi:10.1126/science.abi7178)

67. Eagen KP, Aiden EL, Kornberg RD. 2017 Polycomb-mediated chromatin loops revealed by a subkilobase-resolution chromatin interaction map. Proc Natl Acad Sci U S A 114, 8764–8769. (doi:10.1073/pnas.1701291114)

68. Levo M, Raimundo J, Bing XY, Sisco Z, Batut PJ, Ryabichko S, Gregor T, Levine MS. 2022 Transcriptional coupling of distant regulatory genes in living embryos. Nature 605, 754–760. (doi:10.1038/s41586-022-04680-7)

69. Fujioka M, Mistry H, Schedl P, Jaynes JB. 2016 Determinants of Chromosome Architecture: Insulator Pairing in cis and in trans. PLoS Genet 12, e1005889. (doi:10.1371/journal.pgen.1005889)

70. Georgiev P, Kozycina M. 1996 Interaction between mutations in the suppressor of Hairy wing and modifier of mdg4 genes of Drosophila melanogaster affecting the phenotype of gypsy-induced mutations. Genetics 142, 425–436. (doi:10.1093/genetics/142.2.425)

71. Martin CH, Mayeda CA, Davis CA, Ericsson CL, Knafels JD, Mathog DR, Celniker SE, Lewis EB, Palazzolo MJ. 1995 Complete sequence of the bithorax complex of Drosophila. Proc Natl Acad Sci U S A 92, 8398–8402. (doi:10.1073/pnas.92.18.8398)

