## Supplemental figures for "Boundary Bypass Activity in the *Abdominal-B* Region of the *Drosophila* Bithorax Complex is Position Dependent and Regulated"

### Supplementary information

#### Supplemental Table

Table S1. List of primers for DNA fragments

| Fragment | Primer sequences (5'-3') | Coordinates according to the complete sequence of BX-C in SEQ89E numbering (Martin et al., 1995) | Coordinates according to the FlyBase Drosophila Sequence Coordinates, release=r6.48; species=Dmel |
|---|---|---|---|
| **F8**$^{337}$ | caacgccaaccagcac; aatgtgagttgttagg | 64038-64374 | 3R:16919404..16919068 |
| **F8**$^{165}$ | caacgccaaccagcac; ggtggcgctgcaaggc | 64210-64374 | 3R:16919232..16919068 |
| **F8**$^{209}$ | actttaaatttccacattcc; aatgtgagttgttagg | 64038-64246 | 3R:16919404..16919196 |
| **F7** | gatttcaagctgtgtggcggggg; atgtcggcaattcggattcccgg | 83399-84504 | 3R:16900043..16899468 |
| **F7**$^{dHS1}$ | aagagcgaggtagaatgtcgc; aattccaatcaaaccatcaaac | 84115-84356 | 3R:16899326..16899085 |
| The sequences of the fragments containing multimerized binding sites | | | |
| **Pita**$^{\times 5}$ | *aagctt*(*Hind*III)gatctttagccaagacgcgaacccgaatccgaaactttagccaagacgcgaacccgaatccgaaactttagccaagacgcgaacccgaatccgaaacttagccaagacgcgaacccgaatccgaaactttagccaagacgcgaacccgaatccgaagatctaatatc*gaattc*(*Eco*RI) | | |
| **CTCF**$^{\times 4}$ | *actagt*(*Spe*I)gctgcagcgccacctggccttggagatcctgcagcgccacctggccttggagatcctgcagcgccacctggccttggagatctccaaggccaggtggcgctgcagccccgggctgcag*gaattc*(*Eco*RI) | | |
| **gypsy** | *gaatt*c(*Eco*RI)gatcgcaaaaacattgcatattttcggcaaagtaaaattttgttgcataccttatcaaaaaataagtgctgcatacttttagagaaaccaaataattttttattgcatacccgtttttaataaaatacattgcatacccctcttttaataaaaaatattgcatactttgacgaaacaaattttcgttgcatacccaataaaagattattatattgcatacccgtttttaataaaatacattgcatacccctcttttaataaaaaatattgcatacgttgacgaaacaaattttcgttgcatacccaataaaagattattatattgcatacctttcttgcataccatttagccgat(*Mun*I)*caattg* | | |

**Supplemental Figures**

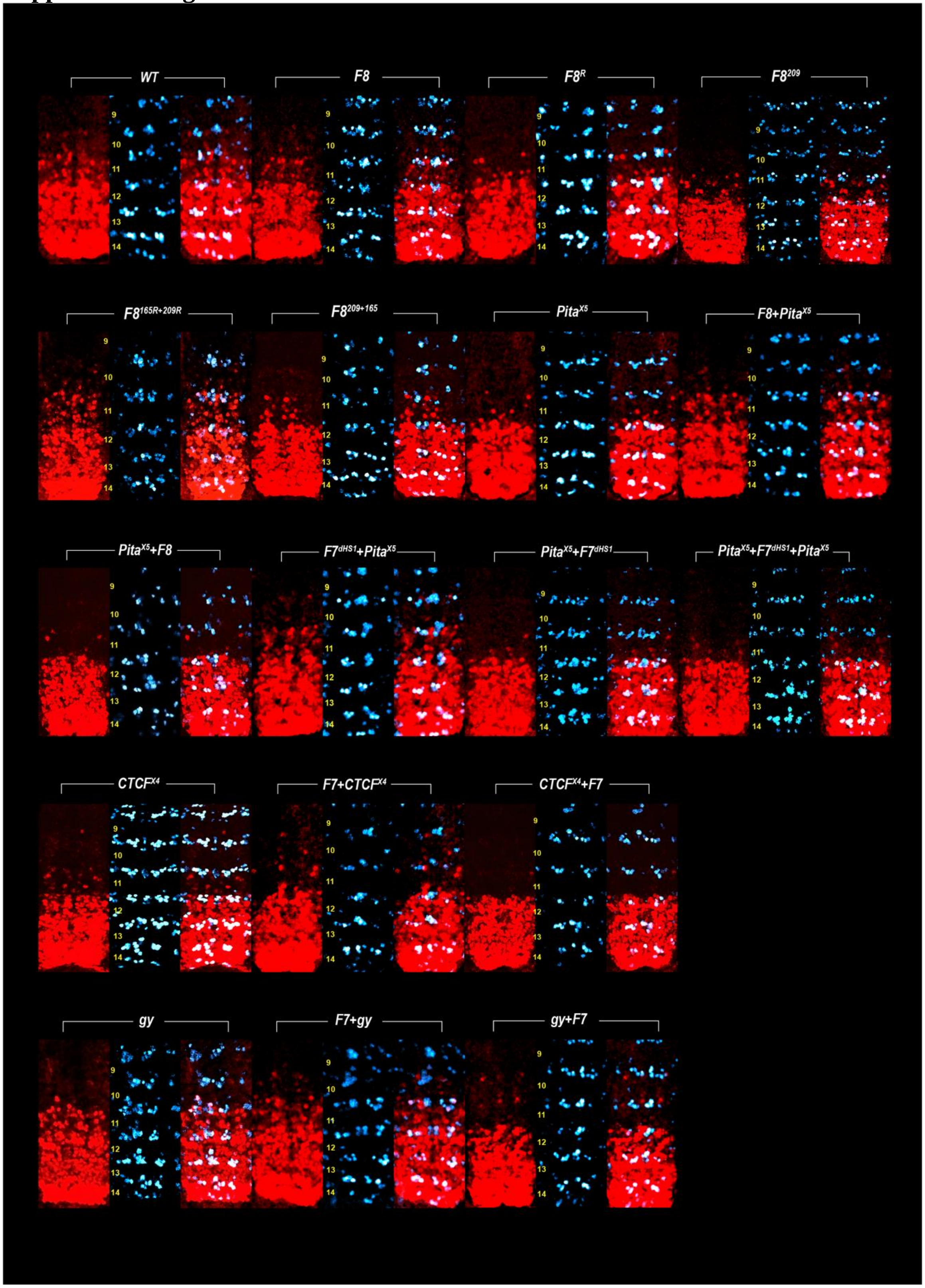


**Fig. S1**. *Abd-B* expression in *Fab-7* replacement embryos. Each panel shows a confocal image of the embryonic CNS from stage 14 embryos stained with antibodies to Abd-B (red) and Engrailed (En, blue). En is used to mark parasegment borders, which are numbered from 9. to 14 on the left side of the panels. Genotype is as indicated.

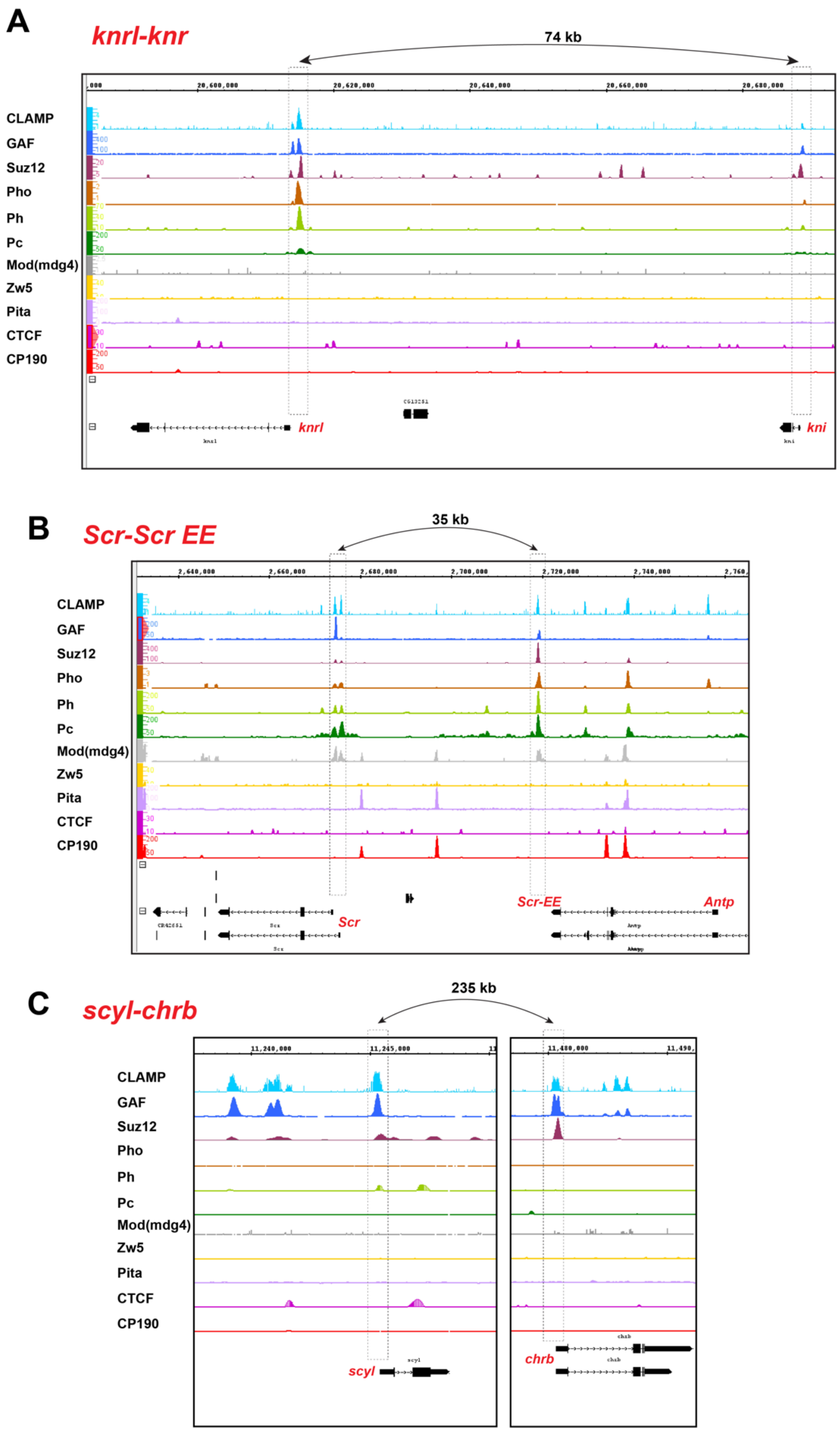
A
knrl-knr
74 kb
20,600,000
20,620,000
20,640,000
20,660,000
20,680,000
CLAMP
GAF
Suz12
Pho
Ph
Pc
Mod(mdg4)
Zw5
Pita
CTCF
CP190
knrl
kni
B
Scr-Scr EE
35 kb
2,640,000
2,660,000
2,680,000
2,700,000
2,720,000
2,740,000
Scr
Scr-EE
Antp
C
scyl-chrb
235 kb
11,240,000
11,245,000
11,480,000
scyl
chrb

**Fig. S2**. Summary results of ChIP-seq data for the binding of for the binding of Suz12, Ph, Pc 1, CTCF, CP190, Zw5, Pita, Elba3, Insv, GAF, CLAMP proteins in knrl-knr (A), the *Scr-ScrEE.* (B) and *scyl-chrb* (C) regions (modENCODE http://www.modencode.org/). The arrows indicate which sequence are linked to each other in each locus. Note that the *SCR EE* maps close to the 5' end of lncRNA, *CR44931*. Abreviations: *knrl*: *knirps-like*; *knr*: *knirps; Scr*: *Sex combs reduced*; *Scr EE*: *Sex combs reduced enhancer*; *scyr*: *scylla*; *chrb*: *charybde*.

List of datasets of proteins

tissue antibody GEO/ENCODE link

PHO E6-18h PHO https://www.ncbi.nlm.nih.gov/geo/query/acc.cgi?acc=GSM1479972

PH E6-18h PH https://www.ncbi.nlm.nih.gov/geo/query/acc.cgi?acc=GSM2211686

Pc E6-18h PC https://www.ncbi.nlm.nih.gov/geo/query/acc.cgi?acc=GSM1479973

E(z) third instar larval imaginal discs and brains E(z)
https://www.ncbi.nlm.nih.gov/geo/query/acc.cgi?acc=GSM2734944

CP190 E5-13h CP190 https://www.ncbi.nlm.nih.gov/geo/query/acc.cgi?acc=GSM1481701

CTCF and Pita as in Maksimenko et al.,2015

Development: doi:10.1242/dev.199827: Supplementary information

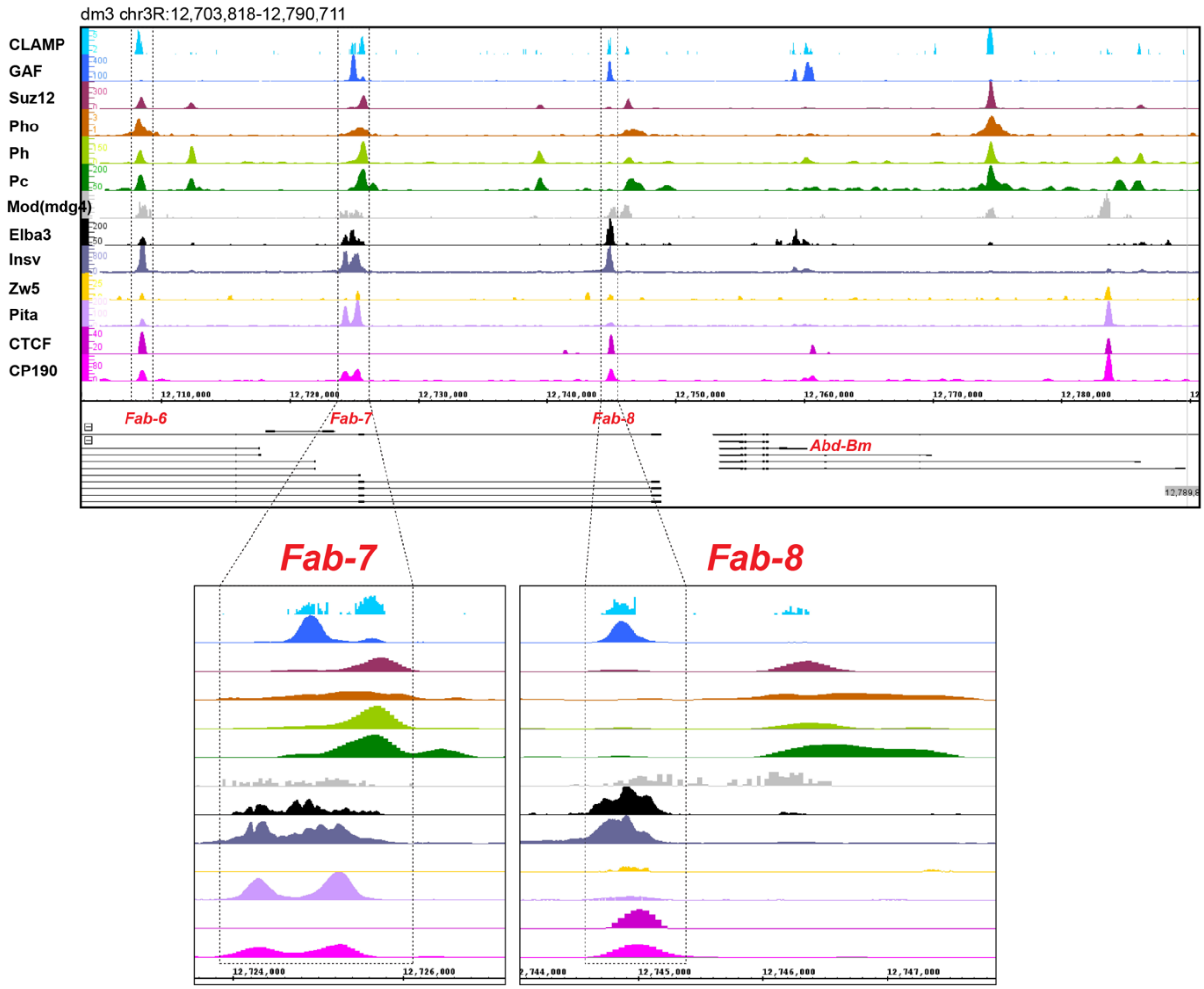

**Fig. S3.** Summary results of ChIP-seq data for the binding of for the binding of Suz12, Ph, Pc 1, CTCF, CP190, Zw5, Pita, Elba3, Insv, GAF, CLAMP proteins in the *Abd-B* region of BX-C (modENCODE http://www.modencode.org/).

List of datasets of proteins

tissue antibody GEO/ENCODE link

PHO E6-18h PHO https://www.ncbi.nlm.nih.gov/geo/query/acc.cgi?acc=GSM1479972

PH E6-18h PH https://www.ncbi.nlm.nih.gov/geo/query/acc.cgi?acc=GSM2211686

Pc E6-18h PC https://www.ncbi.nlm.nih.gov/geo/query/acc.cgi?acc=GSM1479973

E(z) third instar larval imaginal discs and brains E(z) https://www.ncbi.nlm.nih.gov/geo/query/acc.cgi?acc=GSM2734944

CP190 E5-13h CP190 https://www.ncbi.nlm.nih.gov/geo/query/acc.cgi?acc=GSM1481701

CTCF and Pita as in Maksimenko et al.,2015

Development: doi:10.1242/dev.199827: Supplementary information
